# Mixing genome annotation methods in a comparative analysis inflates the apparent number of lineage-specific genes

**DOI:** 10.1101/2022.01.13.476251

**Authors:** Caroline M. Weisman, Andrew W. Murray, Sean R. Eddy

## Abstract

Comparisons of genomes of different species are used to identify lineage-specific genes, those genes that appear unique to one species or clade. Lineage-specific genes are often thought to represent genetic novelty that underlies unique adaptations. Identification of these genes depends not only on genome sequences, but also on inferred gene annotations. Comparative analyses typically use available genomes that have been annotated using different methods, increasing the risk that orthologous DNA sequences may be erroneously annotated as a gene in one species but not another, appearing lineage-specific as a result. To evaluate the impact of such “annotation heterogeneity,” we identified four clades of species with sequenced genomes with more than one publicly available gene annotation, allowing us to compare the number of lineage-specific genes inferred when differing annotation methods are used to those resulting when annotation method is uniform across the clade. In these case studies, annotation heterogeneity increases the apparent number of lineage-specific genes by up to 15-fold, suggesting that annotation heterogeneity is a substantial source of potential artifact.

## Introduction

Comparing the genome sequences of different organisms can yield inferences about the genetic basis of the biological differences between them. One such analysis aims to identify genes unique to a particular monophyletic group. Such genes, called “orphan genes” when restricted to one species and “lineage-specific” or “taxonomically-restricted” when restricted to a clade of several species, are interesting from the perspective of genetic and evolutionary novelty. For example, they have been proposed to underlie lineage-specific structural and functional innovations, and to be novel genes that have emerged from noncoding DNA [1–5].

Lineage-specific genes are typically identified by searching for homologs in outgroup species: genes for which homologs cannot be found are considered lineage-specific. Such analyses typically begin not with raw genome sequences, but with particular “annotations” of them: inferences about what genes they encode. Often, only genes included in these annotations are considered in the homology search [2, 6, 7].

Previous work has recognized two ways in which errors in genome annotations could produce spurious lineage-specific genes. A real gene could be annotated in the focal lineage, but its homologs incorrectly unannotated in outgroups [8–10]. Conversely, a non-genic sequence could be incorrectly annotated as a gene in the lineage, but correctly omitted in outgroups [11]. Such errors could occur even when all genomes in an analysis are consistently annotated by the same annotation methodology, but the potential for error is expected to increase if genomes are annotated by different methods, which use different criteria in determining which sequences are genic. Because comparative analyses typically depend on publicly available genomes whose annotations come from different authors and sources, such “annotation heterogeneity” is common [12–16]. Many gene annotation methods are in wide use, including custom pipelines at large bioinformatics data providers (NCBI [17], Ensembl [18]), hand-curated model organism annotation (Flybase [19], Wormbase [20]), crowd-sourced annotation (VectorBase [21]), and various software packages (Maker [22], PASA [23]), used independently or in combination, with custom parameters chosen by individual researchers.

Here we evaluate the impact of annotation heterogeneity on inferred numbers of lineage-specific protein-coding genes. We identify four clades of species with available genome sequences for which multiple different annotations are publicly available. These enable us to conduct case studies in which we compare the number of lineage-specific genes when all species are annotated with the same method (“uniform annotations”) to when they are annotated with different methods (“heterogeneous annotation”). We find that annotation heterogeneity consistently and substantially increases the inferred number of lineage-specific genes. This effect is strongest when all species within the lineage are annotated with one method and all outgroup species with a different one. Our results suggest that annotation heterogeneity can produce many spurious lineage-specific genes, potentially a majority of those found in a study.

## Results

### Identification of clades of sequenced genomes with annotations from two methods

To directly compare lineage-specific genes found using uniform annotations and heterogeneous annotations, we manually searched the literature and bioinformatic databases for species groups in which all species were annotated with the same method, and, additionally, the same assembly of each species had been independently annotated with some other method. We used existing annotations from a variety of standard sources instead of generating our own to make results maximally representative of real studies. We identified four groups of five species: cichlids, primates, bats, and rodents (Table 1, Supplemental Table 1). For cichlids and primates, all five species were annotated with the same two methods, whereas for bats and rodents, one method was applied to all five species, and the other available annotation was from three different methods, with each species being annotated by one of the three. Each of these four groups is less than approximately 60 My old.

**Table 1:**
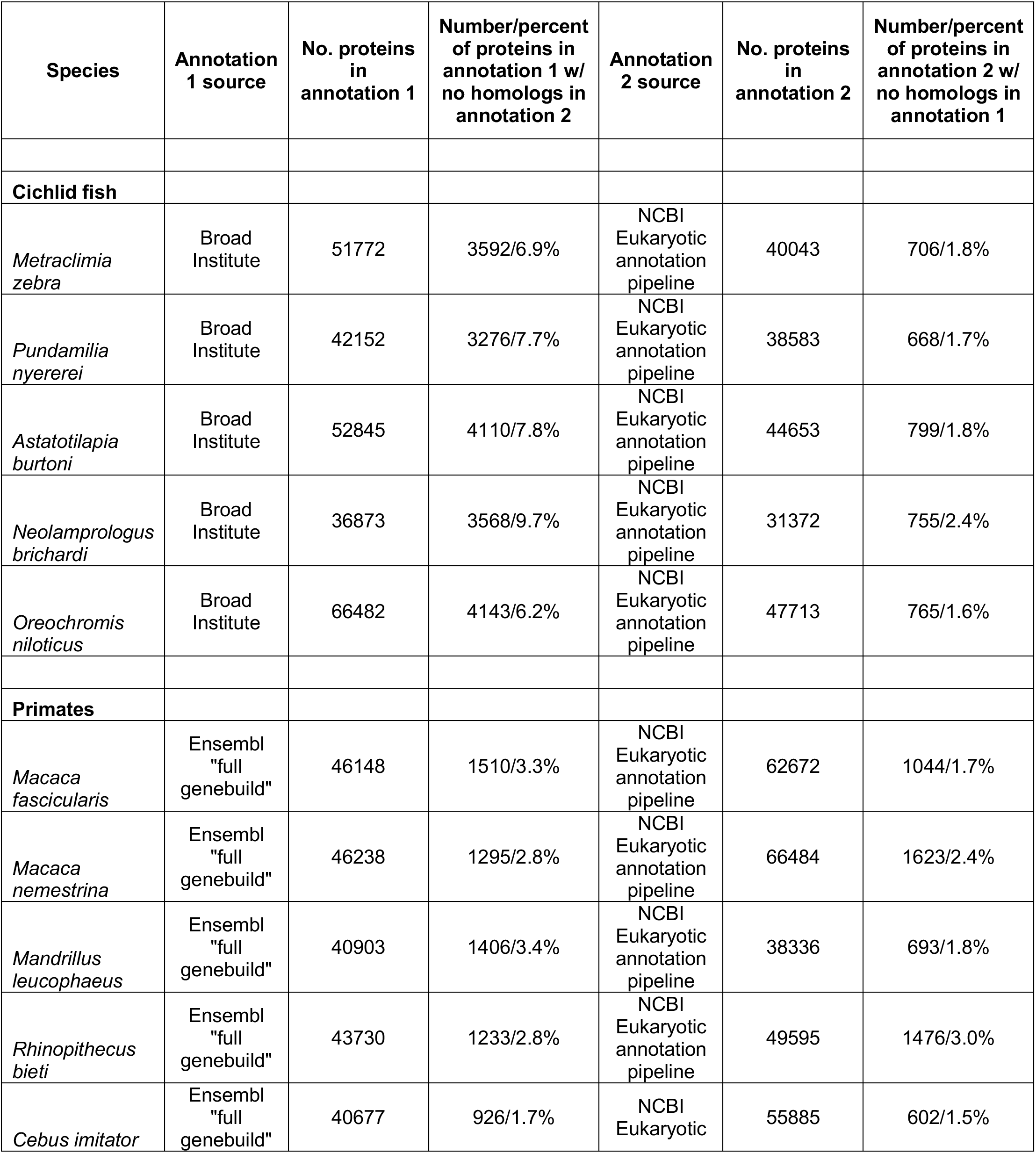
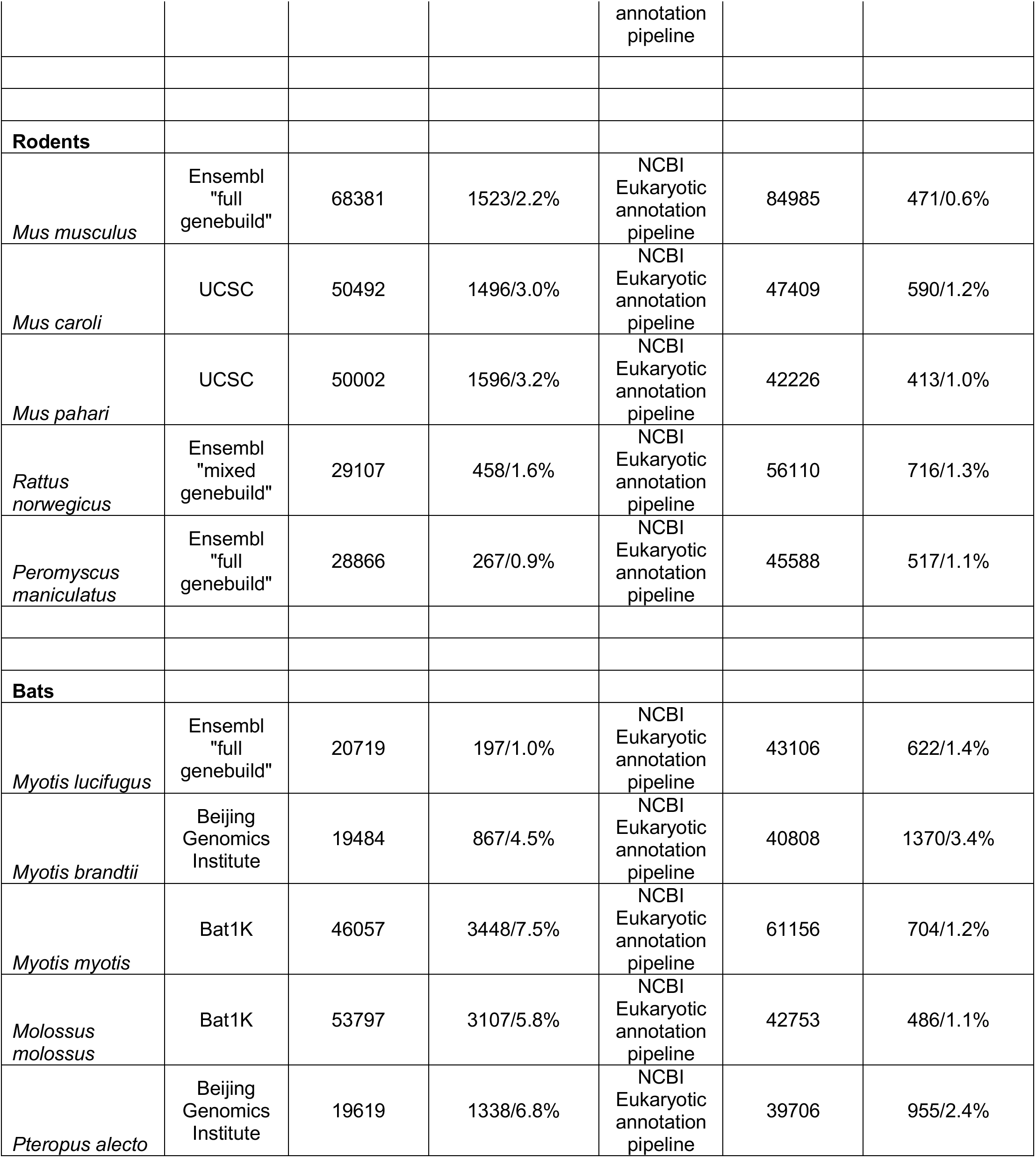
Genome annotations used in this study. Brief description of annotation source, number of genes in the annotation, and the number and percentage of genes in each annotation with no significant homologs found by a BLASTP search in the other annotation for the given species are listed. Note that, where large differences in the number of proteins included in a pair of annotations occurs, this is often due in part to one annotation including a larger number of different isoforms of the same locus, all or many of which may have significant similarity to the same protein(s) in the other annotation.

### Different annotations of the same genome have many proteins unique to each method

Spurious lineage-specific genes may result from annotation heterogeneity when different annotation methods differentially annotate homologous sequences. Spurious lineage-specific genes may also result from such erroneous differential annotation even when a single annotation method is used, as sequence differences between the species may alter a given method’s determination regarding genic status. To get a sense of how many spurious lineage-specific protein-coding genes annotation heterogeneity *per se* can produce, we compared two protein annotations of the same species to identify proteins appearing to be unique to one of the annotations. Because the underlying genome sequences are identical, any such apparently unique proteins must be spurious, due only to annotation heterogeneity.

To mimic a typical analysis, for each species’ two annotations, we used BLASTP [24] for all proteins in one annotation to see if a significantly similar (E<0.001) homolog was present in the other annotation. Between 0.6% and 9.7% of proteins in one annotation had no significantly similar sequence in the other (Table 1). Of the 40 (20 pairs) annotations, 19 had over 1000 proteins without a significant homolog in the other annotation. In an extreme case of the cichlid *Astatotilapia burtonii*, one annotation (Broad Institute) found 4110 genes that had no significant similarities in the other (NCBI eukaryotic annotation pipeline), and 799 proteins in the NCBI annotation lacked significant similarities in the Broad annotation. These substantial differences between two annotations of one genome illustrate the potential for spurious lineage-specific genes in comparisons of different genomes.

### Different patterns of annotation heterogeneity may differently affect the inferred number of lineage-specific genes

When different annotation methods are used for species within an analysis, different patterns in which those methods are arranged on the species topology are possible. These different patterns may differently affect the number of spurious lineage-specific genes produced by annotation heterogeneity. In particular, because a gene is called as “lineage-specific” if no significant homologs are found in any species outside the lineage, we expected that the number of spurious lineage-specific genes would be positively related to the overall degree of difference between the lineage and outgroup annotations.

We considered three such patterns. In the first, one annotation method is used for all ingroup species (in the lineage, the gray boxes in the figures), and a different method for all outgroup species (outside the lineage); we refer to this as “phyletic” annotation (Figure 1). In the second, one method is used for all ingroup species, but a mixture of methods is used for the outgroup species; we refer to this as “semi-phyletic” annotation (Figure 2). In the third, a mixture of methods is used for both the ingroup species and the outgroup species; we refer to this as “unpatterned” annotation (Figure 3). We used our four clades to create case studies for each pattern.

**Figure 1:**
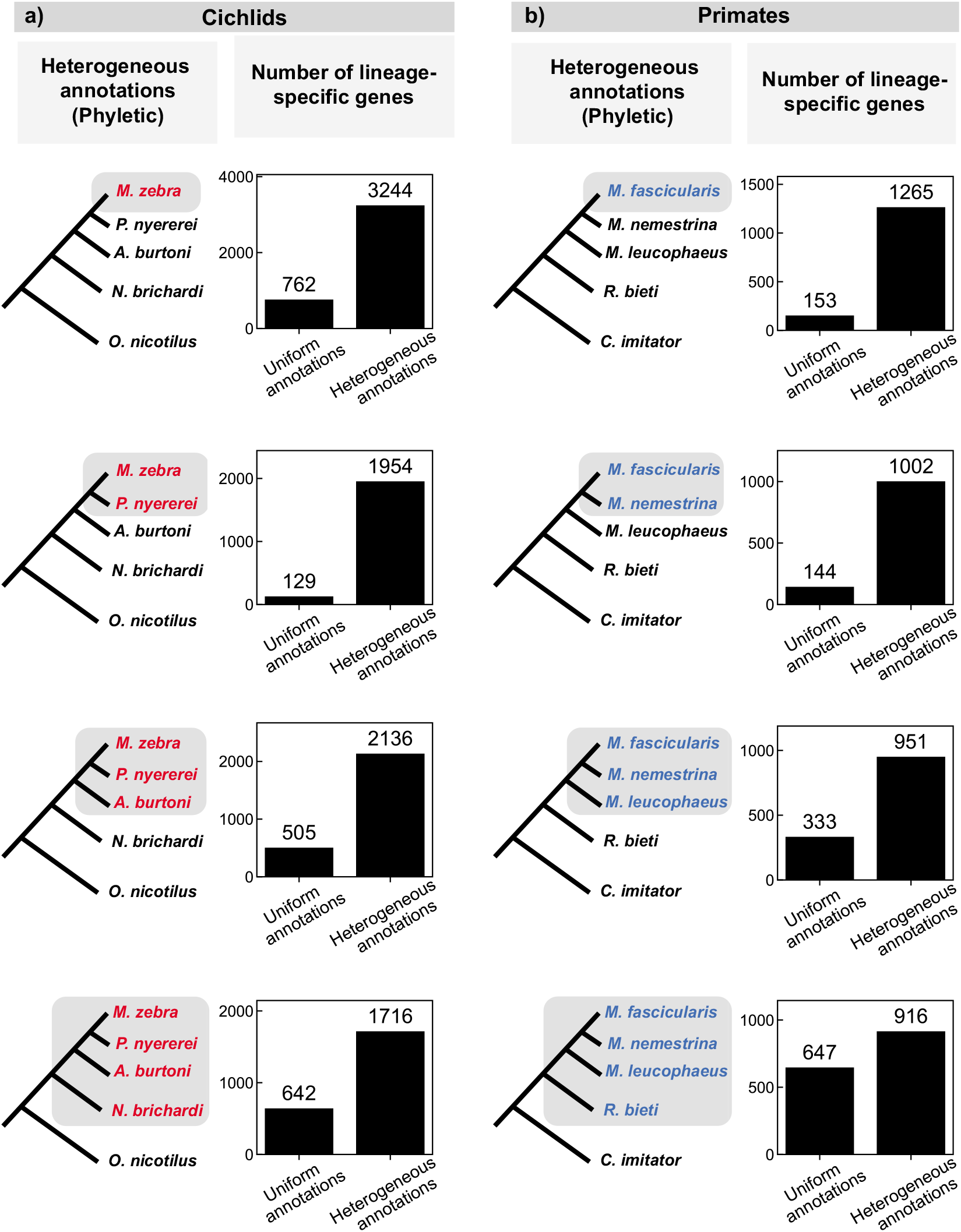
Comparison of the number of lineage-specific genes found using uniform and heterogeneous (phyletic) annotations in a) cichlids and b) primates. The species tree on the left indicates the lineage under consideration (grey shading); different text colors indicate different annotation sources in the heterogeneous annotation analysis (black, NCBI; red, research group at the Broad Institute; blue, Ensembl). A depiction of the uniform annotation pattern, in which all annotations are from NCBI (black), is not shown. Bar graphs indicate the number of genes that appear specific to the lineage shaded on the species tree to the left using either uniform or heterogeneous annotations.

**Figure 2:**
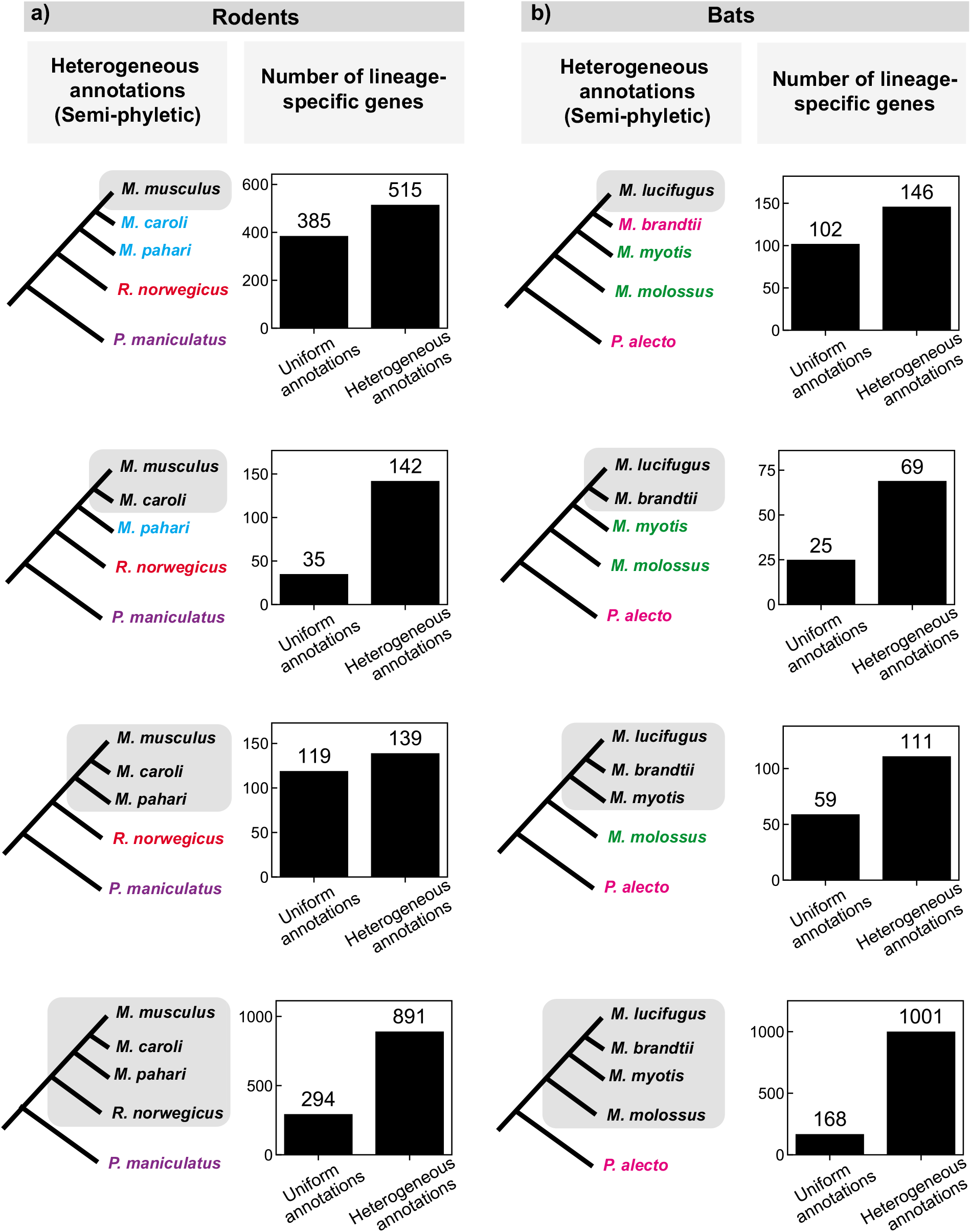
Comparison of the number of lineage-specific genes found using uniform and heterogeneous (semi-phyletic) annotations in a) rodents and b) bats. The species tree on the left indicates the lineage under consideration (grey shading); different text colors indicate different annotation sources in the heterogeneous annotation analysis (black, NCBI; blue, UCSC; red, Ensembl “mixed genebuild”; purple, Ensembl “full genebuild”; green, Bat1k; pink, Beijing Genomics Institute). A depiction of the uniform annotation pattern, in which all annotations are from NCBI (black), is not shown. Bar graphs indicate the number of genes that appear specific to the lineage shaded on the species tree to the left using either uniform or heterogeneous annotations.

**Figure 3:**
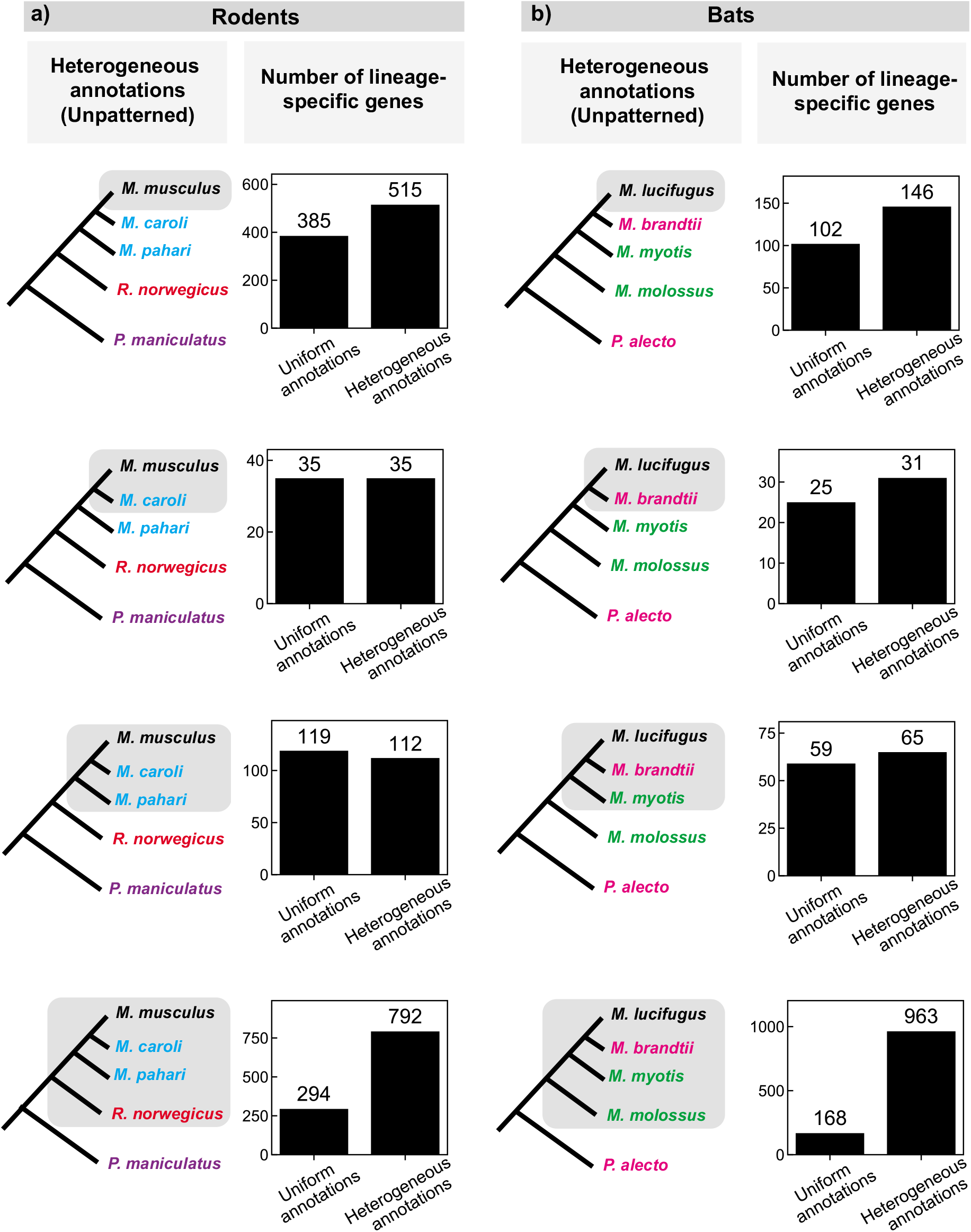
Comparison of the number of lineage-specific genes found using uniform and heterogeneous (unpatterned) annotations in a) rodents and b) bats. The species tree on the left indicates the lineage under consideration (grey shading); different text colors indicate different annotation sources in the heterogeneous annotation analysis (black, NCBI; blue, UCSC; red, Ensembl “mixed genebuild”; purple, Ensembl “full genebuild”; green, Bat1k; pink, Beijing Genomics Institute). A depiction of the uniform annotation pattern, in which all annotations are from NCBI (black), is not shown. Bar graphs indicate the number of genes that appear specific to the lineage shaded on the species tree to the left using either uniform or heterogeneous annotations.

The differences in annotation methods between ingroups and outgroups are largest for phyletic annotation, intermediate for semi-phyletic annotation, and smallest for unpatterned annotation; we expected the number of spurious lineage-specific genes to scale accordingly.

### Annotating a lineage with one method and outgroups with a different method greatly increases the apparent number of lineage-specific genes

Phyletic annotation occurs in at least two scenarios. Studies that newly sequence a lineage often use their own method to annotate that lineage, and may then compare it to outgroup annotations from another single source (e.g. Ensembl). Additionally, studies using existing annotations may encounter a correlation between taxon and annotation method because genome sequencing groups (with their annotation teams) often select species taxonomically (e.g. studies of particular taxa, sequencing consortia/database initiatives for particular taxa) [25, 26].

We tested the impact of phyletic annotation on the apparent number of lineage-specific genes on two groups of species, where the same genome assembly for every species had been annotated by the same two methods: five cichlids, annotated both by the Broad Institute and NCBI; and five primates, annotated both by Ensembl and NCBI (Supplemental Table 1).

For each tree of five species, we exploited the ladder-like topology (Figure 1) of the tree to perform four analyses, comparing each of the four monophyletic groups including the focal species to the remaining outgroups. For each lineage that included the focal species, we conducted a typical analysis of lineage-specific genes by identifying genes in the focal species that have a significantly similar homolog in the deepest rooted member of the ingroup (and thus are “present” in that clade), but lack significant similarity to any protein in any outgroup species in a BLASTP search (Methods). We compared the number of lineage-specific genes found when all species (both ingroups and outgroups) were annotated with the same method to the number found when the annotations for all outgroup species were switched to the other method in a “phyletic” annotation pattern (Figure 1).

Heterogeneous annotation consistently caused a large increase of hundreds to thousands of apparent lineage-specific genes, typically about a 4-fold (ranging from 1.4-fold to 15-fold) difference relative to uniform annotation. In all but one of the eight cases in Figure 1, the increase is more than 2-fold, suggesting that the majority of lineage-specific genes inferred in heterogeneous annotations are artifacts of the heterogeneity.

### Annotating a lineage with one method and outgroups with a mixture of other methods increases the apparent number of lineage-specific genes

Examples of what we call “semi-phyletic” annotation, where the ingroup is annotated with one method and outgroups with a mixture of methods, are common in the literature on lineage-specific genes [12, 13, 26–31]. This can occur in scenarios similar to phyletic annotation, but where outgroup annotations are available from a mixture of sources (e.g. a combination of Ensembl and NCBI).We created case studies of semi-phyletic annotation using groups of species for which every species had been annotated both by the same method and by one of a mix of other methods: five rodents and five bats (Supplemental Table 1). We repeated the procedure described for phyletic annotation above to compare the number of lineage-specific genes in semi-phyletic annotations to those in uniform annotations (Figure 2).

Semi-phyletic annotation heterogeneity caused a smaller but still substantial increase in the number of apparent lineage-specific genes in all lineages in both groups (Figure 2). The magnitude of this effect ranged from 20 to 833 additional lineage-specific genes, corresponding to 1.2-fold to 6-fold increases.

### Annotating species with a mixture of methods without taxonomic bias increases the apparent number of lineage-specific genes

Examples of what we call “unpatterned” annotation, where the annotation method varies within the ingroup as well as the outgroup, are also common in the literature [15, 16, 27, 32–35]. This occurs when studies use existing available annotations for the desired species, which may come from a variety of sources. We created case studies of unpatterned annotation using the same rodent and bat species we used for semi-phyletic annotation (Figure 2), with the difference that we always compared the uniform annotations to the full set of mixed annotations (Figure 3) to produce unpatterned annotation heterogeneity.

Unpatterned annotation heterogeneity usually caused an increase in apparent lineage-specific genes (Figure 3), though the effect was smaller than for phyletic or semi-phyletic annotations. Two cases showed equal numbers or slight decreases, and the other six cases showed increases of 1.1-fold to 5.7-fold; the largest increases were in the cases with a single outgroup species.

### As expected, six-frame translation homology searches dramatically reduce the apparent number of lineage-specific genes

A homology search in which the query protein is compared directly to a six-frame translation of the target genome does not rely on an annotation of the target species, and so should reduce this source of spurious lineage-specific genes. Such translated searches have previously been shown to reduce the inferred number of lineage-specific genes [8, 9]. In agreement with these expectations, we find that, for all of the lineages described above (depicted in Figures 1–3), a search for the focal species’ proteins against six-frame translations of all comparator species genomes dramatically reduces the number of lineage-specific genes: to below the number inferred with uniform annotations, and often to less than one hundred (Supplemental Table 2).

## Discussion

We used six case studies to ask if varying the annotation method across species in a comparative analysis (“annotation heterogeneity”) alters the apparent number of lineage-specific genes. We found that switching from uniform to heterogeneous annotations consistently increased the number of genes that were classified as lineage-specific, with increases ranging from tens to thousands of genes, corresponding to increases of up to 15-fold. The largest increases were seen when one annotation method was used for all the ingroup species and another was used for all the outgroup species (“phyletic annotation”). The smallest increases were seen when a mixture of annotation methods were used in both ingroup and outgroup species. Our case studies consist of trees of five species; mixtures of annotations in larger numbers of outgroup and ingroup species may reduce the artifact.

Annotation heterogeneity is common in comparative studies. Our results suggest that the numbers of lineage-specific genes found in these studies may be inflated, especially in “phyletic annotation” cases, and where the number of species compared is small. Annotation heterogeneity may also have consequences that we do not explore here, like producing spurious lineage-specific losses.

Recent work from us and others has shown that homology detection failure, in which homology searches fail to detect homologs that are actually present in outgroups, can also produce spurious lineage-specific genes [36, 37]. Previous studies have noted a surprisingly large number of “young” lineage-specific genes found in recently evolved clades [15], which, compared to older lineage-specific genes, are less readily explained by homology detection failure, which is minimized at short evolutionary distances. The results here are all for young (<60 My old) clades, showing that annotation heterogeneity can be a significant source of spurious lineage-specific genes in young clades.

In accordance with previous results, we show that annotation heterogeneity artifacts can be reduced by performing homology searches of six-frame translated genomic DNA sequence in search of unannotated homologs in target species. This approach has caveats. At short evolutionary distances, a sequence may be sufficiently similar for successful detection in such a search without having the same coding status as the query; for example, a truly de novo originated gene is expected to have significant nucleotide similarity to a homologous noncoding locus in close outgroup species. This approach also still relies on an accurate annotation of the focal species.

When annotation methods disagree, which is correct? Our results do not address this, only demonstrating a consequence of this disagreement. Even homogeneous annotations are imperfect. Of particular concern, methods in general rely on features (homology to known genes, length, expression level, codon optimization) that seem likely to be absent or weaker in newly evolved (*de novo*) genes, and so may fail to identify these genes. We consider annotation accuracy primarily accountable to experimental data. Testing transcription, translation, and function in all species in question is of ultimate importance in accurately identifying lineage-specific genes. In light of our results, we suggest more emphasis on these metrics. In the meantime, the true number of lineage-specific genes remains difficult to ascertain, but better understanding sources of spurious ones helps us constrain it.

## Author Contributions

Conceptualization, C.W.; Formal Analysis, C.W.; Investigation, C.W.; Writing – Original Draft: C.W.; Writing – Review & Editing: C.W., A.W.M, S.R.E; Supervision, A.W.M, S.R.E; Funding Acquisition, S.R.E.

## Declaration of interests

The authors declare no competing interests.

## Methods

### Identifying lineage-specific proteins

For each species group, we defined a protein as specific to a particular lineage if a search using BLASTP [24] version 6.2.0 had no similar protein at a significance threshold of E=0.001 in the annotation of any species that was an outgroup to that lineage. We did not require that a protein be present in all members of the lineage to be specific to that lineage: a protein was defined as specific to a lineage based on the most distant species in which it was detected. For example, if a protein in *M. musculus* was detected only in *R. norvegicus*, it was defined as specific to that lineage; if a gene in *M. musculus* was detected in *M. caroli*, *M. pahari*, *and R. norvegicus*, it was also defined as specific to that same lineage. If a protein was found in the earliest-branching member of the species group, it was considered “conserved” and so not counted as any kind of lineage-specific gene. This way of classifying lineage-specificity coheres with standard practice [6].

For the six-frame translated searches, we first generated a six-frame translation of the genome assembly of each species using the ‘esl-translate’ command in the hmmer easel package, and then used it as the target database in a BLASTP search, as described in the previous paragraph.

## Supplemental Information

Supplemental Table 1: Sources, brief descriptions, and links to protein annotations and genome assemblies used in this study.

Supplemental Table 2: Results of six-frame translation homology searches. Numbers in the table indicate the inferred number of genes specific to the indicated lineage (corresponding to the four lineages depicted in Figures 1–3) in each of the described taxa.

## Data availability

All raw results summarized in Figures 1–3 are available at https://github.com/caraweisman/Annotation_homology.

## Notes

### Competing Interest Statement

The authors have declared no competing interest.

